# Comparison and critical assessment of single-cell Hi-C protocols

**DOI:** 10.1101/2022.05.08.491066

**Authors:** M. Gridina, A. Taskina, T. Lagunov, A. Nurislamov, T. Kulikova, A. Krasikova, V Fishman

**Author notes:** correspondence should be sent to Veniamin Fishman. equal contribution (M. Gridina, A. Taskina).

## Abstract

Advances in single-cell sequencing technologies make it possible to study the genome architecture in single cells. The rapid growth of the field has been fueled by the development of innovative single-cell Hi-C protocols. However, the protocols vary considerably in their efficiency, bias, scale and costs, and their relative advantages for different applications are unclear.

Here, we compare the two most commonly used single-cell Hi-C protocols. We use long-read sequencing to analyze molecular products of the Hi-C assay and show that whole-genome amplification step results in increased number of artifacts, larger coverage biases, and increased amount of noise compared to PCR-based amplification. Our comparison provides guidance for researchers studying chromatin architecture in individual cells.

## Introduction

Chromatin architecture plays an important role in genome biology. Hi-C is one of the most common techniques employed to study chromatin organization in the nucleus. Multiple modifications of Hi-C protocol were developed to study genome architecture (Kempfer and Pombo, 2020), yet most of them require large amount of input material and therefore can not be applied to study individual cells.

Single cell analysis is essential when pure subpopulations cannot be isolated or when obtaining large amount of cells is challenging. Typical examples of such rare cell populations are oocytes and early embryonic cells (Díaz et al., 2018; Flyamer et al., 2017; Zhang et al., 2020). Recently, several single-cell Hi-C (scHi-C) protocols have been proposed to address this challenge. The key difference between these protocols is a strategy of DNA amplification. Whereas in the protocols based on method by (Nagano et al., 2013) library amplification is accomplished by a routine PCR, protocols developed by (Flyamer et al., 2017) employ phi29 polymerase with the strand-displacement activity for the whole-genome amplification (WGA) of proximity ligation products. Due to different amplification strategy, WGA-protocols do not allow enrichment of proximity ligation products; in addition, these protocols differ in DNA fragmentation methods (see Table 1 to review key differences of scHi-C methods

| Article | Key protocol features |
| --- | --- |
| Flyamer et al., 2017 | Multiple displacement amplification (MDA); Sonication |
| Nagano et al., 2017 | PCR-based; Tagmentation of ligated DNA, Biotin enrichment |
| Collombet et al., 2020 | PCR-based; Tagmentation of ligated DNA; Biotin enrichment |
| Stevens et al., 2017 | PCR-based, AluI digestion of ligated DNA; Biotin enrichment |
| Tan et al., 2018 | Multiplex end-tagging amplification of ligated DNA |
| Ke et al., 2017 | PCR-based; Biotin enrichment |
| Ulianov et al., 2021 | Multiple displacement amplification (MDA); Sonication |

Here, we compared two basic subtypes of scHi-C protocol suitable for the analysis of rare cell populations: WGA-and PCR-based. We found that PCR-based amplification generates more uniform coverage and reduced number of artifacts compared to phi29-based amplification. Using long-read sequencing of phi29-derived products, we showed that phi29-amplification results in circular DNA overamplification and template switching, which explains many of the observed artifacts.

## Methods

### Nuclei isolation and formaldehyde fixation

Chicken oocytes with a diameter from 1 to 2 mm were dissected from the ovary and placed in individual drops of cooled “5:1” medium (83 mM KCl, 17 mM NaCl, 6.5 mM Na2HPO4, 3.5 mM KH2PO4, 1 mM MgCl2, 1 mM DTT, pH 7.2). Nuclei were isolated as previously published (Krasikova et al., 2012), washed with “5:1” medium and transferred for fixation into a dish containing 2% formaldehyde in PBS for 15 min at room temperature. Formaldehyde was quenched with 0.125 M glycine during 15 min at room temperature. The nuclei integrity was estimated under Olympus stereomicroscope. The Hi-C protocol was proceeded immediately.

### Single-nucleus Hi-C

Two Hi-C protocols were used in this study:

The first one - WGA-based. We applied the previously described protocol by Flyamer et al., 2017 with minor modifications regarding the NGS libraries preparation, which were prepared with KAPA HyperPrep kit.

Two samples were prepared for ONT sequencing according to the manufacturer manual.

The second one was PCR-based.

The main differences from the previous one were using :

- UltraPure™ Low Melting Point Agarose (Thermo Fisher Scientific) to avoid nuclei loss;
- biotine-dCTP for the enrichment of the ligation products;
- AluI for fragmentation of DNA for NGS libraries preparation;
- 25 cycles of PCR.

Individual fixed nuclei were transferred into wells of microplate with a 9μL ice-cold lysis buffer for 20 min. Nuclei were washed with NEB3.1 + 0.6% SDS, and moved to 0.2 ml PCR tubes with the 3μL NEB3.1 + 0.6% SDS. 10 μL fresh prepared 1.2% Low Melting Point Agarose were added into each tube with control under the a stereomicroscope. After solidification 10 μL NEB3.1 + 0.6% SDS were added and samples were incubated 1 hrs at 37°C. 10 μL 6% Tritоn X-100 were used for SDS quenching and chromatin fragmentation was performed with 25U DpnII (New England Biolabs) at 37°C for overnight. The digested chromatin ends labeling was performed by 5 U Klenow fragment in the presence of biotin-15-dCTP at 22°C for 4 hrs followed by ligation at 16°C for overnight. Crosslinks were reversed by incubating at 65°C overnight and Low Melting Point Agarose were digested by Agarase (Thermo Fisher Scientific) 1 hr at 42°C. DNA was fragmented by AluI 1 hr at 37°C and NGS libraries were prepared with KAPA HyperPrep kit according to the manufacturer manual with some modifications. The volumes of end repair, adapter ligation reactions and PCR were reduced by 5, 5 and 2 times, respectively. The adapters were diluted to 300 nM. There was biotin pull-down after the adapter ligation step. 25 cycles of PCR were performed.

### Bioinformatic analysis

Illumina libraries processing was performed using Juicer software as described in (Fishman et al., 2019). ONT-data was processed using Pore-C-Snakemake version 0.2.0.

Rings ratio (RR) was obtained by dividing total number of the aligned read base pairs (Aligned) by total length of unique reference regions (Unique) covered by this read and subtracting obtained ratio from 1: RR = 1 -Aligned / Unique. Rings ratio equal to zero indicates that each alignment block reported for the read is unique and does not overlap with other alignment blocks, whereas rings ratio close to one indicates that alignment blocks are highly overlapping. Randomized controls of reads ratio were obtained by randomly permuting positions of reads.

### Data availability

Sequencing data is available via NCBI SRA, PRJNA834620.

## Results and discussion

### PCR-based amplification generates better single-cell Hi-C libraries compared to WGA-based protocols

We prepared single-cell Hi-C libraries from nuclei of chicken oocytes using two different library preparation methods using two different library preparation methods: WGA-and PCR-based (Fig. 1). We used oocytes because they represent typical cells that can not be analyzed using conventional Hi-C protocols. Although we note that studying chromatin organization in these cells is important, we limit the current study to the comparison of Hi-C protocols.

**Figure 1.**
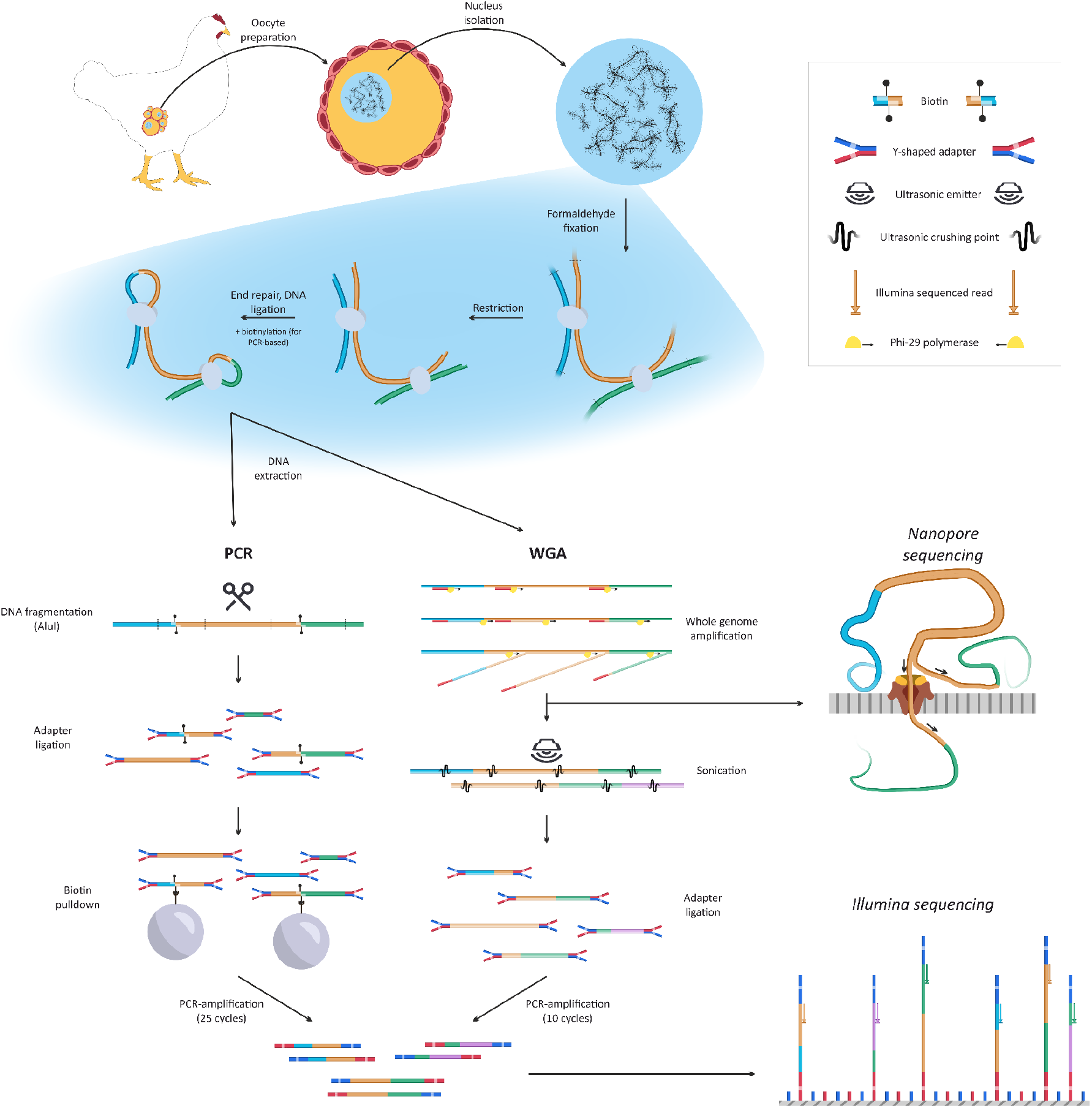
Two protocols of single-cell Hi-C library preparation benchmarked in this study. Both methods include nucleus isolation, nucleus fixation, chromatin restriction, DNA ends repair and ligation. PCR-based protocol continues with AluI-digestion, biotin pull-down, adapter ligation, amplification and Illumina sequencing. WGA-based protocol includes phi29-amplification followed either by direct ONT-sequencing or sonication, adapter ligation, PCR amplification and Illumina sequencing.

We sequenced nine libraries prepared with the WGA-based protocol and 15 libraries prepared using PCR-based protocol. The details of these protocols are provided in the “Methods” section, but here we briefly review the key differences between them. Both protocols begin with nucleus isolation, formaldehyde fixation, and in-chromatin digestion with DpnII enzyme (Fig. 1). The PCR-based protocol continues with DNA ends biotinylation and ligation, DNA fragmentation, biotin pull-down which allows enrichment of ligation products, adapter ligation, and PCR-amplification. For WGA-based protocol, we amplified DNA after the ligation step using phi29 enzyme, which has a strand-displacement activity, fragmented the obtained library using sonication, and continued with adapter ligation, DNA amplification, and Illumina sequencing.

Using sequencing data, we accessed several key statistics of Hi-C libraries. First, we computed the number of intra-fragment read pairs, representing DNA fragments that failed to ligate. We found that WGA-based protocol results in 4-5 times more intra-fragment reads than PCR-based protocol (Fig. 2, A). We assumed that this difference is because in the WGA-based protocol, there is no enrichment of proximity ligation products. In accord with the increase of intra-fragment reads, for WGA-based libraries, we observed more reads in FR-orientation, which often (although not always - Gridina et al., 2021) inversely correlate with the number of proximity ligation products (Fig. 2, A). Finally, we note that both WGA-and PCR-based libraries are PCR-amplified after DNA fragmentation and adapter ligation steps, and aimed to estimate library complexity after this final round of amplification. We found that coverage distribution was more uniform for PCR-based libraries, whereas in WGA-based libraries several loci showed unexpectedly high sequencing coverage (Fig. 2, A).

**Figure 2.**
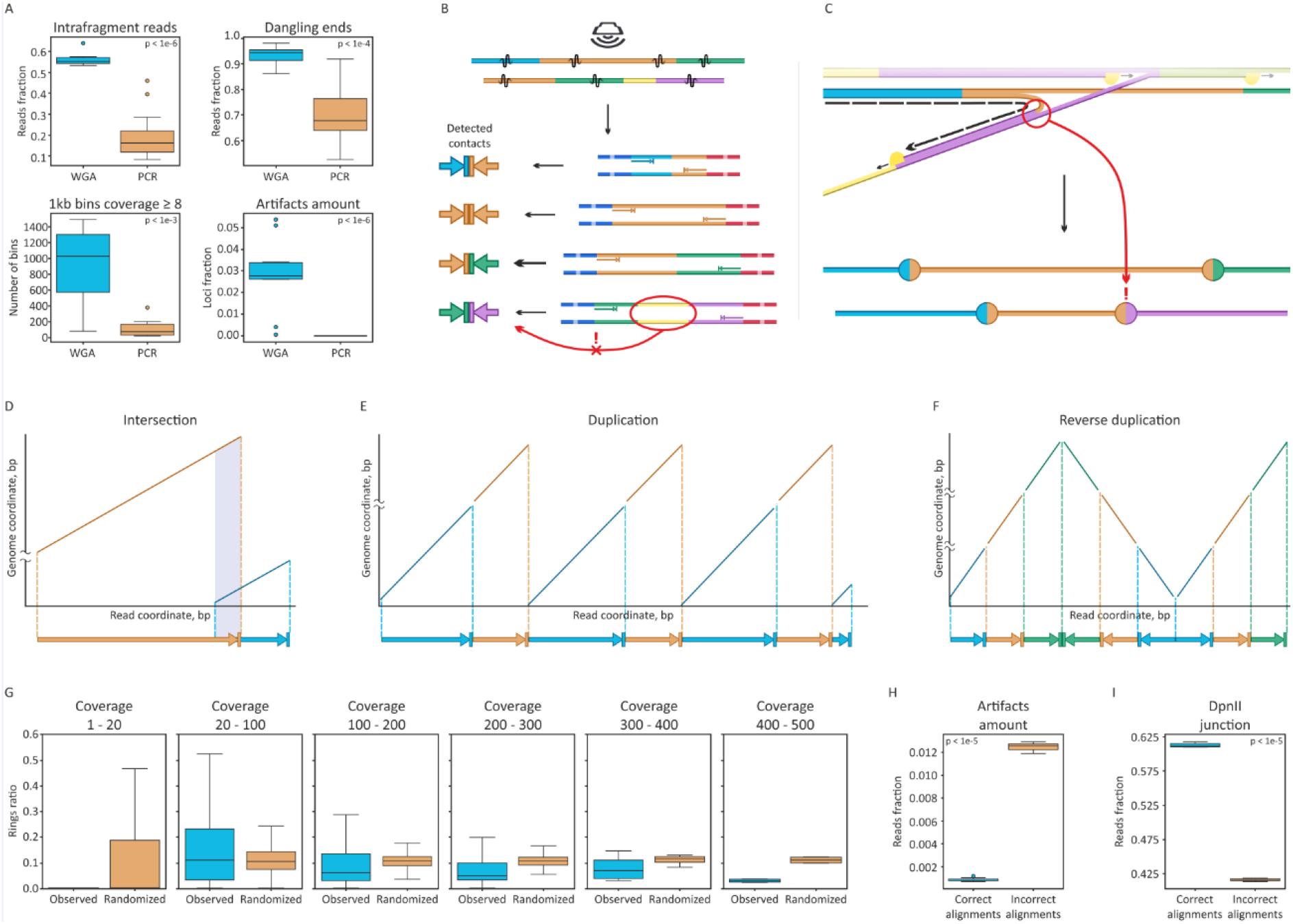
Comparison of scHi-C protocols. A. Quality metrics for PCR-and WGA-based protocols. B and C. Illustration of two hypotheses explaining artifacts observed in scHi-C data. D-F. Representations of different alignment types observed in long-read sequencing data. Graphs represent real alignment results observed in the data. G. Interdependence of genomic coverage of loci and rings ratio of the reads overlapping them. Briefly, rings ratio equal to zero indicates that each alignment block reported for the read is unique and does not overlap with other alignment blocks, whereas rings ratio close to one indicates that alignment blocks are highly overlapping. H. Fraction of artifacts shown for reads characterized by consistent and inconsistent alignments. I. Fraction of artifacts shown for reads with or without DpnII junctions. For A, I and H, samples represent individual oocyte nuclei, p-values obtained using Mann-Whitney U-test.

Interestingly, some restriction fragments within these highly covered regions displayed numerous interactions, whereas we expected not more than eight interactions to be observed for each restriction fragment, i.e. four (number of chromatids in meiotic oocyte) * 2 (ends of each fragment) (Galitsyna and Gelfand, 2021). We defined restriction fragments detected in more than eight different chimeric DNA fragments as artifacts and counted the number of artifacts for each of the obtained libraries. To compare this statistics across samples, we downsampled all datasets to 20 000 read pairs to remove a factor of sequencing depth. This analysis shows that the number of artifacts was substantially higher for WGA-based libraries than for PCR-based (Fig. 2, A).

### Artifacts in WGA-based libraries arise from circular fragments overamplification and template switching

We proposed two alternative hypotheses explaining artifacts of the WGA-based protocol (Fig. 2, B and C).

First, random fragmentation of identical long molecules may produce short fragments containing different pairs of restriction fragments on their ends (Fig. 2, B). These different short fragments are interpreted as independent Hi-C-contacts, although they represent WGA duplicates of the single long proximity ligation product. This leads to an overestimation of contact counts and loci proximity. Second, phi29 polymerase may switch templates during amplification (Lasken and Stockwell, 2007), generating new DNA junctions which are erroneously interpreted as products of proximity ligation (Fig. 2, C).

To score the importance of these two sources of artifacts and to explain why some loci display abnormally high coverage after WGA, we employed Oxford Nanopore long-read technology (ONT). We split all material obtained after WGA into two aliquots, one sequenced directly using ONT, and another subjected to sonication, adapter ligation, and Illumina short-read sequencing (Fig. 1). This allowed us to directly compare Hi-C statistics obtained for short-and long-read sequencing.

We examined ONT-read alignments within the regions displaying abnormally high coverage. Typical ONT-reads contain junctions of several genomic fragments, which often concur with DpnII cut sites. However, in regions displaying high coverage, we observed reads containing multiple repeats of individual DpnII restriction fragments (Fig. 2, E and F). Frequently, but not always, the order of the fragments within the read was unchanged, i.e. the same chain of segments was repeated multiple times (Fig. 2, E). However, we also observed cases when the chain of fragments was reordered, i.e. a single read contained genomic segments A, B and C ordered as A-B-C-C-B-A (Fig. 2, F).

We interpret these repetitive chains of fragments as products obtained by amplification of circular templates, generated during the proximity ligation step. Quantitative analysis also confirmed that the portion of circular products significantly increases at regions with high genomic coverage (Fig. 2, G). We assumed that reordering of segments arises from polymerase template switching, as described in (Lasken and Stockwell, 2007).

To sum up, we concluded that WGA amplification results in unequal reads coverage due to the overamplification of circular products. Template switching during amplification of circular templates produces artifacts that do not represent proximity ligation events.

We next aimed to explore other sources of artifacts in the WGA-derived scHi-C data. For this aim, we explored alignments of the ONT reads, which supported contacts of fragments with more than eight contacts. We found that these reads often contain inconsistent alignment segments, which we defined as alignment segments with sharing some fraction of read sequence (Fig. 2, D) or alignment segments separated by a large unaligned gap. Filtering out overlapping alignments or segments separated by a large (>= 10 bp) unaligned gap significantly reduced the number of artifacts (Fig. 2, H). Moreover, we found that DpnII cut sites, which are expected at Hi-C ligation junctions, coincide with the ends of consistent alignment segments significantly more often than with the ends of inconsistent alignment segments. Filtering our reads without DpnII cut sites near alignment segment junctions also significantly reduced the number of artifacts (Fig. 2, I). Thus, we concluded that erroneous alignment is an important source of artifacts in ONT-derived Hi-C data. Interestingly, analysis of the same WGA library subjected to sonication and Illumina sequencing with the matched total number of contacts showed a substantially lower number of artifacts, indicating that erroneous alignments are mainly an issue of the long-reads mapper.

Finally, we aimed to estimate the portion of artifacts arising at the sonication step. As described previously, random sonication may result in different fragments originating from similar copies of long proximity ligation products (Fig. 2, B). In ONT reads, we were able to count indirect junctions of DNA fragments, i.e. when two DNA fragments are separated by a third fragment. Such indirectly interacting fragments may be considered as directly interacting after sonication and Illumina-sequencing if read length is not sufficient to detect an intervening DNA fragment. We found that exclusion of such indirect contacts reduces the number of artifacts only slightly (from 5,67% to 5,61%), indicating that this source of artifacts is not essential.

## Conclusions

Our data shows that PCR-based scHi-C protocols provide better data quantity and quality compared to WGA-based protocols. The vast majority of artifacts observed in WGA data arise from phi29 polymerase template switching. Random sonication does not add much to the number of the artifacts. Long-read sequencing makes it possible to find products of circular overamplification and distinguish between direct and indirect contacts. However, accurate aligning of long chimeric reads is challenging, and segment alignments must be carefully filtered to avoid artifacts.

## Acknowledgments

This work, including cells isolation and NGS library preparation, was supported by Russian Science Foundation grant #20-64-46021. Illumina sequencing was performed by Skoltech Genomics Core Facility. Long-read sequencing was performed using equipment of the Novosibirsk State University, supported by the Ministry of Education and Science of Russian Federation, grant #2019-0546 (FSUS-2020-0040). Computations were performed on the nodes of HPC cluster of the Institute of Cytology and Genetics, supported by the project No. 121031800061-7).

## Authors contribution

V.F. and A.K. conceived the study. M.G. and A.N. isolated oocytes with help from T.K. and A.K. M.G. prepared and sequenced Hi-C libraries. A.N. performed ONT-sequencing. A.T. performed bioinformatic analysis with help from T.L. and V.F. All authors contributed to manuscript preparation.

## Notes

### Competing Interest Statement

The authors have declared no competing interest.

## Bibliography

Collombet, S., Ranisavljevic, N., Nagano, T., Varnai, C., Shisode, T., Leung, W., Piolot, T., Galupa, R., Borensztein, M., Servant, N., et al. (2020). Parental-to-embryo switch of chromosome organization in early embryogenesis. Nature 580, 142–146. https://doi.org/10.1038/s41586-020-2125-z.

Díaz, N., Kruse, K., Erdmann, T., Staiger, A.M., Ott, G., Lenz, G., and Vaquerizas, J.M. (2018). Chromatin conformation analysis of primary patient tissue using a low input Hi-C method. Nat. Commun. 9, 4938. https://doi.org/10.1038/s41467-018-06961-0.

Fishman, V., Battulin, N., Nuriddinov, M., Maslova, A., Zlotina, A., Strunov, A., Chervyakova, D., Korablev, A., Serov, O., and Krasikova, A. (2019). 3D organization of chicken genome demonstrates evolutionary conservation of topologically associated domains and highlights unique architecture of erythrocytes’ chromatin. Nucleic Acids Res. 47, 648–665. https://doi.org/10.1093/nar/gky1103.

Flyamer, I.M., Gassler, J., Imakaev, M., Brandão, H.B., Ulianov, S.V., Abdennur, N., Razin, S.V., Mirny, L.A., and Tachibana-Konwalski, K. (2017). Single-nucleus Hi-C reveals unique chromatin reorganization at oocyte-to-zygote transition. Nature 544, 110–114. https://doi.org/10.1038/nature21711.

Galitsyna, A.A., and Gelfand, M.S. (2021). Single-cell Hi-C data analysis: safety in numbers. Brief. Bioinform. 22, bbab316. https://doi.org/10.1093/bib/bbab316.

Gridina, M., Mozheiko, E., Valeev, E., Nazarenko, L.P., Lopatkina, M.E., Markova, Z.G., Yablonskaya, M.I., Voinova, V.Y., Shilova, N.V., Lebedev, I.N., et al. (2021). A cookbook for DNase Hi-C. Epigenetics Chromatin 14, 15. https://doi.org/10.1186/s13072-021-00389-5.

Ke, Y., Xu, Y., Chen, X., Feng, S., Liu, Z., Sun, Y., Yao, X., Li, F., Zhu, W., Gao, L., et al. (2017). 3D Chromatin Structures of Mature Gametes and Structural Reprogramming during Mammalian Embryogenesis. Cell 170, 367–381.e20. https://doi.org/10.1016/j.cell.2017.06.029.

Kempfer, R., and Pombo, A. (2020). Methods for mapping 3D chromosome architecture. Nat. Rev. Genet. 21, 207–226. https://doi.org/10.1038/s41576-019-0195-2.

Krasikova, A., Khodyuchenko, T., Maslova, A., and Vasilevskaya, E. (2012). Three-dimensional organisation of RNA-processing machinery in avian growing oocyte nucleus. Chromosome Res. 20, 979–994. https://doi.org/10.1007/s10577-012-9327-7.

Lasken, R.S., and Stockwell, T.B. (2007). Mechanism of chimera formation during the Multiple Displacement Amplification reaction. BMC Biotechnol. 7, 19. https://doi.org/10.1186/1472-6750-7-19.

Nagano, T., Lubling, Y., Stevens, T.J., Schoenfelder, S., Yaffe, E., Dean, W., Laue, E.D., Tanay, A., and Fraser, P. (2013). Single-cell Hi-C reveals cell-to-cell variability in chromosome structure. Nature 502, 59–64. https://doi.org/10.1038/nature12593.

Nagano, T., Lubling, Y., Várnai, C., Dudley, C., Leung, W., Baran, Y., Mendelson Cohen, N., Wingett, S., Fraser, P., and Tanay, A. (2017). Cell-cycle dynamics of chromosomal organization at single-cell resolution. Nature 547, 61–67. https://doi.org/10.1038/nature23001.

Stevens, T.J., Lando, D., Basu, S., Atkinson, L.P., Cao, Y., Lee, S.F., Leeb, M., Wohlfahrt, K.J., Boucher, W., O’Shaughnessy-Kirwan, A., et al. (2017). 3D structures of individual mammalian genomes studied by single-cell Hi-C. Nature 544, 59–64. https://doi.org/10.1038/nature21429.

Tan, L., Xing, D., Chang, C.-H., Li, H., and Xie, X.S. (2018). Three-dimensional genome structures of single diploid human cells. Science 361, 924–928. https://doi.org/10.1126/science.aat5641.

Ulianov, S.V., Zakharova, V.V., Galitsyna, A.A., Kos, P.I., Polovnikov, K.E., Flyamer, I.M., Mikhaleva, E.A., Khrameeva, E.E., Germini, D., Logacheva, M.D., et al. (2021). Order and stochasticity in the folding of individual Drosophila genomes. Nat. Commun. 12, 41. https://doi.org/10.1038/s41467-020-20292-z.

Zhang, K., Wu, D.-Y., Zheng, H., Wang, Y., Sun, Q.-R., Liu, X., Wang, L.-Y., Xiong, W.-J., Wang, Q., Rhodes, J.D.P., et al. (2020). Analysis of Genome Architecture during SCNT Reveals a Role of Cohesin in Impeding Minor ZGA. Mol. Cell 79, 234–250.e9. https://doi.org/10.1016/j.molcel.2020.06.001.

